# Template-driven dynamic functional network connectivity predicts medication response for major depression and bipolar disorders

**DOI:** 10.1101/2024.10.17.618523

**Authors:** Bradley T. Baker, Elizabeth Osuch, Scott Langenecker, Jay Fournier, Jessica Turner, Eric Youngstrom, Vince D. Calhoun

## Abstract

The process of finding reliable treatment for major depression and bipolar disorder can be arduous. The myriad behavioral symptoms presented by patients and resistance to treatment from particular medication classes complicate standard diagnostic and prescription methodologies, often requiring multiple attempted treatments during which symptoms may still be present. Physiological information such as neuroimaging scans may help to alleviate some of the uncertainty surrounding diagnosis and treatment when incorporated into a clinical setting. Changes in functional magnetic resonance imaging show particular promise, as the incorporation of dynamical information may provide insights into physiological changes prior to static, structural changes. In this work, we present a novel method for generating robust and replicable dynamic functional network connectivity (dFNC) features from neuroimaging data using a template of dynamic states derived from a large, non-affected data set. We demonstrate that this template-driven dFNC approach expands on standard dFNC approaches by allowing for the derivation of a continuous state-contribution time series. We demonstrate that the derived biomarkers can support high predictive performance for the identification of medication class and non-responders while also expanding the set of biomarkers available for studying differences in mood disorder medication response.

## 1. INTRODUCTION

It has long been hypothesized that different mood disorders, such as major depressive disorder (MDD) and bipolar disorder (BPD) differ in their neurological basis [1, 2, 3]. Despite these underlying differences, behavioral symptoms for MDD and BPD may appear similar, especially when individuals with BPD are experiencing a depressive phase. This difficulty in distinguishing between diagnosis in clinical practice can lead to prolonged trial treatment periods, where patients may still experience symptoms if the initial diagnosis was incorrect based on similar behavioral symptoms. Treatment resistant depression (TRD) and individual differences in treatment response effectiveness can further complicate matters when certain individuals may exhibit behavioral signs of depression but not respond to standard medication classes [4, 5, 6]. In response to these difficulties, a significant amount of work has been put forward identifying biomarkers that can predict individual responses to specific medications, providing clinicians with tools for more personalized treatment [4, 5, 3, 6, 7]. A further class of machine-learning methodologies utilize these biomarkers to build predictive models which can be actively used in clinical decision support. For a survey of recent (mostly genomic) works performing treatment-response prediction for Major Depression see [8].

Functional magnetic resonance imaging (fMRI) is a neuroimaging technique that measures blood oxygenation level-dependent (BOLD) contrast during a scanning session [9]. Biomarkers derived from fMRI have proven useful in differentiating between MDD and BPD [10, 11, 12] as well as in predicting responses to specific medications [5, 3, 7]. Resting-state fMRI, in particular, has revealed intrinsic connectivity networks (ICNs) such as the striatum precuneus [10] and the default mode and fronto-parietal networks [12], which provide insights for further research into mood disorders and treatment responses. One of the potential advantages of functional MRI over other modalities is that the BOLD response is measured over a period of time, allowing for the study of neurological dynamics such as dynamic functional network connectivity (dFNC). While a majority of prior works have focused on utilizing genomic factors or other static measures for determining medication response[8], the dynamic aspects of functional imaging also show immense promise for this challenging problem.

Multiple prior studies [7, 13] have studied medication responses using spatially constrained independent component analysis (scICA) with the Neruomark template [14]. Salman et al. utilized seven domain-specific resting-state networks from Neuromark, selected through sequential forward selection (SFS) with a kernel support vector machine (SVM) model. Across three preceding studies [3, 5, 15], the SVM model achieved about 90% accuracy when non-responders were excluded. In a more recent study[13], version 2.0 of the Neuromark template[16] was utilized, which incorporates resting-state networks across multiple spatial resolutions [17]. This study demonstrates that ICNs from this multi-scale template serve as effective biomarkers for predicting medication response. In this work, when spatial maps were combined with static functional network connectivity (sFNC) figures, an AUC of 0.93 was achieved, with non-responders included. While promising, these works utilize derivatives that collapse the dynamic nature of the fMRI signal into static measures.

In this work, we substantially expand on the application of spatially constrained ICA to the problem of treatment response prediction, introducing new biomarkers which fully exploit the dynamic nature of the fMRI signal. First, we apply standard dynamic functional network connectivity (dFNC) using both the Neuromark version 1.0 and 2.0 templates. Furthermore, we go beyond standard dFNC by utilizing spatially constrained ICA with a dFNC state template, derived from a large data set with 100k plus non-affected individuals [17]. This template-driven dynamic functional network connectivity (TD-dFNC) allows us to derive a set of states which align spatially with the known non-affected states, but which may appear to demonstrate different dynamics within the affected population. We compare the performance of both of these dynamic biomarkers as input to a kernel-SVM classifier, providing a direct comparison to previous studies [7, 13]. We demonstrate that the dynamics derived with dFNC and the TD-dFNC provide comparable or improved performance to the static derivatives utilized in previous studies, further highlighting the promise of fully utilizing dynamic fMRI derivatives for medication-response prediction.

## 2. METHODS

In this section, we describe the methodologies utilized in this work. First, we discuss the data set utilized for this study. We then provide background on Neuromark ICA and the version 1.0 and 2.0 templates which were utilized. We then provide an overview of dynamic functional network connectivity. We then continue by presenting our novel template-driven dFNC method. At last, we discuss the kernel-SVM and feature selection methods utilized to gauge medication-class prediction performance using each biomarker type.

### 2.1. Data Set

As our primary aim is to compare novel dynamic biomarkers with previous work on medication response, we utilize a fMRI data set collected at Western University [3, 7] as the primary data source for our study. The participants, aged 16 to 27, showed no significant age differences between groups (p = 0.1492). Diagnoses were made using either the Structured Clinical Interview for DSM Disorders-IV (SCID-IV) or the Diagnostic Interview for Genetic Studies (DIGS) and confirmed by clinical psychiatric assessments. A consensus between SCID-IV/DIGS and clinical diagnoses was required for inclusion in the study. The diagnostic groups included 147 subjects in total: 33 controls, 35 with bipolar disorder (BD) type-I, 67 with major depressive disorder (MDD), and 12 with unknown diagnoses. Participants with conflicting diagnoses or with first-degree relatives suffering from mental illness were assigned to the ”unknown” group. Chart review was utilized to determine the medication class, with the primary goal being sustained euthymia lasting at least six months. Medications were categorized as either antidepressants (AD) or mood stabilizers (MS), which included lithium, lamotrigine, carbamazepine, and divalproex sodium. Based on medication class, the 147 subjects were grouped as follows: 33 controls, 47 responding to AD, 45 responding to MS, 8 non-responders, and 14 who remitted without medication.

MRI data were collected using a 3.0T Siemens Verio MRI scanner with a 32-channel phased-array head coil at the Lawson Health Research Institute. Resting-state fMRI scans were obtained using a gradient-echo, echo-planar imaging (EPI) sequence with the following parameters: repetition time (TR) = 2000 ms, echo time (TE) = 30 ms, 40 axial slices of 3 mm thickness, no parallel acceleration, a flip angle of 90°, field of view (FOV) = 240 × 240 mm, and matrix size = 80 × 80. The resting-state scan lasted approximately 8 minutes, and 164 brain volumes were collected. Data preprocessing and quality control were conducted using the SPM software, following the steps outlined in [7].

### 2.2. Neuromark ICA

First, to extract intrinsic connectivity networks (ICNs) from resting-state fMRI data, we used the Neuromark multi-scale template (Neuromark fMRI 2.1, http://trendscenter.org/data), which comprises 105 networks extracted across multiple spatial resolutions [17]. These ICNs were derived from over 20 different studies using a group multi-scale ICA approach [16], covering eight distinct model orders to capture networks at various spatial scales. Higher model orders generally reflect higher spatial resolution, while lower orders integrate features from broader brain regions. This multi-scale approach enables the modeling of a diverse set of ICNs. Of the 900 extracted components, 105 were retained after manual selection based on criteria such as peak activation in gray matter, minimal overlap with vascular or ventricular structures, and low similarity to artifacts [17]. These components span six functional domains: visual (VI), cerebellar (CB), temporal (TP), subcortical (SC), somatomotor (SM), and higher cognitive (HC).

For comparison, we also included 53 ICNs from the original Neuromark template, following the analysis in [18, 7]. These ICNs are grouped into seven functional domains: sub-cortical (SC), auditory (AD), sensorimotor (SM), visual (VI), executive control (CO), default mode (DM), and cerebellar (CB).

We applied spatially constrained ICA (scICA) using the GIFT fMRI toolbox [14] to extract ICNs. Additionally, we computed the functional network connectivity (FNC) matrix for each participant using Pearson correlation between the time courses of the 105 ICNs, producing a 105 × 105 FNC matrix for each subject. For comparison, we repeated the scICA+FNC analysis using the 53 ICNs from the original Neuromark template. Following the procedure in [7], these spatial maps and FNC matrices were used as features for subsequent prediction tasks.

### 2.3. Dynamic Functional Network Connectivity

Dynamic functional network connectivity (dFNC) [19] extends static functional connectivity by utilizing a sliding window analysis on derived ICA time courses. In standard dFNC, Sliding Window Pearson Correlation (SWPC) is computed with a window size of *W*, so that if *T* is the number of TRs in the data, *T − W* windows are obtained. Each of these windows has the same dimension as the static FNC matrix (105×105 for Neuromark 2.0, and 53×53 for Neuromark 1.0). These dFNC windows can be utilized as learning features (see Table 1).

**Table 1.** The mean and standard deviation performance of each feature type in both unimodal and multimodal scenarios. Each feature type was computed independently on the Western data set[3], and for spatial maps, the components identified in [13] were utilized for the sake of direct comparison. For each method, the mean AUC and F1 scores and their standard deviation over 5-fold cross-validation and 10 random model initializations are shown. Scores above 0.93 for AUC and 0.89 for F1 are highlighted. The version numbers refer to v1.0 and v2.0 of the Neuromark template discussed above.

| Features Used | AUC | F1 |
| --- | --- | --- |
| v1.0 Spatial Maps[7] | 0.9032 ± 0.02998 | 0.8141 ± 0.02309 |
| v1.0 Spatial Maps + sFNC[7] | 0.9175 ± 0.02925 | 0.8840 ± 0.02812 |
| v1.0 dFNC Windows | 0.9040 ± 0.03175 | **0.9010 ± 0.02566** |
| v1.0 dFNC States | 0.8809 ± 0.05189 | 0.8245 ± 0.02356 |
| v1.0 Spatial Maps + dFNC Windows | 0.9053 ± 0.05086 | 0.8809 ± 0.02432 |
| v1.0 Spatial Maps + dFNC States | 0.9156 ± 0.02778 | 0.8853 ± 0.01993 |
| v2.0 Spatial Maps[13] | **0.9368 ± 0.0002** | 0.8375 ± 0.004116 |
| v2.0 Spatial Maps + sFNC[13] | 0.9229 ± 0.02455 | 0.8670 ± 0.03254 |
| v2.0 dFNC Windows | 0.9100 ± 0.03185 | **0.8992 ± 0.02202** |
| v2.0 dFNC States | 0.9148 ± 0.05126 | 0.8561 ± 0.03065 |
| v2.0 Spatial Maps + dFNC Windows | 0.9179 ± 0.02855 | **0.8904 ± 0.02286** |
| v2.0 Spatial Maps + dFNC States | 0.8797 ± 0.05565 | 0.8853 ± 0.02286 |
| v2.0 TD-dFNC Time Series | 0.9169 ± 0.02898 | **0.8968 ± 0.01064** |
| v2.0 TD-dFNC States | **0.9361 ± 0.0015** | 0.8867 ± 0.02496 |
| v2.0 Spatial Maps + TD-dFNC Time Series | 0.9103 ± 0.03220 | 0.8878 ± 0.02374 |
| v2.0 Spatial Maps + TD-dFNC States | **0.9300 ± 0.01827** | **0.8904 ± 0.02643** |

Post-processing on dFNC takes the form of a two-stage K-Means clustering: first, a set of exemplar windows are computed by measuring the standard deviation in connectivity over all participant windows and selecting the windows within each participant, which represent local maxima of the standard deviation time-course. These exemplar windows are then utilized to determine the optimal *K* number of states by measuring the silhouette score over repeated clustering for different values of *K*. The elbow criterion is utilized to determine an optimal *K* based on the exchange of variance explained and diminishing returns by adding more clusters. Once the number of states is determined, all windows can then be clustered together so that each participant’s windows are assigned to a particular state. The average states across participants and further derivatives, such as the number of state transitions and mean dwell time, can then be utilized for statistical comparison between groups.

In this work, we detrend, despite, and apply a low-pass filter at 0.15 HZ to the ICA time-courses prior to computing the K-Means clustering. We utilize a window size of 30. For Neuromark 1.0, we found *K* = 4 to be the optimal number of states, and for Neuromark 2.0, we found *K* = 5 to be optimal; however, we reduced the Neuromark 2.0 analysis to *K* = 4 to allow for a more direct comparison with the 1.0 template. We utilize K-Means clustering as implemented in the GIFT toolbox using correlation distance, K++ initialization, and 150 max iterations. No clusterings required 150 iterations to converge. In Table 1, we refer to the windows derived prior to the clustering and the “states” which, for each participant, are the averaged windows associated with a particular state following the K-Means clustering.

### 2.4. Template-Driven dFNC

Standard dFNC requires utilizing sliding window-pearson correlation followed by a K-means clustering in order to derive a set of dynamic FNC “states” and, for each participant, an assignment of each of their sliding windows to one of these states. One of the limitations of this approach for smaller data sets is that the derived states may be heavily biased towards the particular data set for which they were derived, thus limiting the generalizability of the discovered biomarkers to other participants in other studies. The Neuromark ICA framework circumvents this for standard ICA on fMRI data by deriving a large template of spatial maps from a consortium of 100k individuals across multiple studies [14]. Recent work has also invested in the creation of a similar dFNC “template” [20] from the same large data set of 100k participants.

Formally, we can optimize the joint objective function from spatially constrained ICA [14] given as:

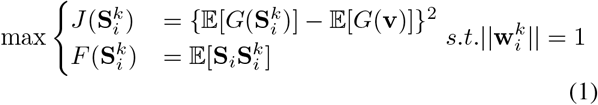

where **S**_*i*_ denotes the *i*th state from the template and 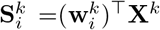 is the estimated state from the *k*th subject, derived from *X*^*k*^ the participant’s windows computed via sliding-window pearson correlation. The vector (*v*) is a Gaussian variable with zero mean and unit variance, and *G* is a non-quadratic nonlinear function applied to this variable. The function *J* is the negentropy of the derived state, and the function *F* represents the correspondence between the derived state and the template state.

This approach, which we call template-driven dFNC (TD-DFNC), exploits the same spatially constrained ICA trick utilized for Neuromark ICA but is now applied to dynamic variables in the form of dFNC windows. In contrast to standard dFNC, this analysis produces a continuous “contribution time-course” for each derived state rather than a binary transition vector. Both derivatives can be used for classification, and in Table 1, we refer to this time series and the fully derived states.

### 2.5. Medication-Class Identification

To classify patient medication responses, we utilized a support vector machine (SVM), following the methodology described in [7]. SVMs are a popular choice for classification tasks, relying on a linear classifier that maximizes the separation between classes. For tasks that require more complexity, we can introduce a nonlinear decision boundary using a kernel function. In our analysis, we employed a Riemannian kernel based on the principal angle between subspaces (PABS) distance metric. This metric is calculated using the singular values derived from the matrix product **U**^*⊤*^ **V**, where **U** and **V** represent spatial maps in voxel-by-component space. The nonlinear kernel is formulated as *K*(**U, V**) = tanh(*γS*(**U, V**)), with the scaling parameter *γ* set to 1. We repeated this process for each subject to generate a comprehensive *N × N* kernel matrix.

As features for classification, we exhaustively explore a number of derivatives both from previous work and novel derivatives from this one. Column 1 in Table 1 lists the full sets of features. From version 1 of Neuromark [14], we utilized ICA spatial maps, static functional network connectivity, dFNC windows, and dFNC states. From version 2 of Neuromark [16], we utilized ICA spatial maps, static functional network connectivity, dFNC windows, and dFNC states, the TD-dFNC contribution time-series and the TD-dFNC states. We also used combinations of features as denoted by the “+” sign in column 1 of Table 1. To compute the kernel for multimodal features, we concatenate the singular values from each domain prior to computing the kernel as denoted above.

For all classification tasks, we perform 5-fold cross-validation and randomly initialize the model 10 times to obtain 50 performance metrics. We then provide the mean validation AUC and F1 in columns 2 and 3 of Table 1, with standard deviations computed over all folds and initializations.

### 3. RESULTS AND DISCUSSION

In this section, we describe the new results presented in this work and discuss their implications.

First, in Table 1, we present the classification results obtained for all of the features derived in this work. Looking at the Neuromark version 1.0 results, we can see that all of the dFNC-based derivatives perform comparably to the spatial map and sFNC derivatives from previous work, with only the version 1.0 dFNC states performing markedly worse than previous approaches using that template. When utilized alongside the spatial maps, the dFNC states provide comparable performance to the best v1.0 model using the sFNC. This suggests that with the current approach and when using the version 1.0 template, the dynamic information from the dFNC windows or their associated states does not help to better predict treatment response.

When looking at the results using the version 2.0 template, we see that the spatial maps from previous work [13] provide better performance than the dFNC windows or states by themselves. Indeed, combining the spatial maps with the dFNC windows does not significantly increase performance. (i) TD-dFNC state 1 (j) TD-dFNC state 2 (k) TD-dFNC state 3 (l) TD-dFNC state 4(m) TD-dFNC state 5(n) TD-dFNC state 6

Interestingly, however, the states derived from TD-dFNC provide comparable performance to the spatial maps, even when the spatial maps are not included as additional features. The other TD-dFNC derivative, the contribution time series, performs comparably to the dFNC derivatives. This performance suggests that the rich information within the TD-dFNC template can be transferred to the problem of medication prediction quite successfully. The dynamic information also seems to hold useful information, although not to the level or stability of the states or spatial maps.

In figure 1, we plot the 4 states derived from standard dFNC using the Neuromark 1.0 (a-d) and 2.0 (e-h) templates, respectively. In figure 1, we also plot the 6 states derived from TD-dFNC (i-n). In each cell of these plots, the Pearson correlation values between each pair of ICNs are provided, with all plots using a uniform scale of -1 to 1.

**Fig. 1.**
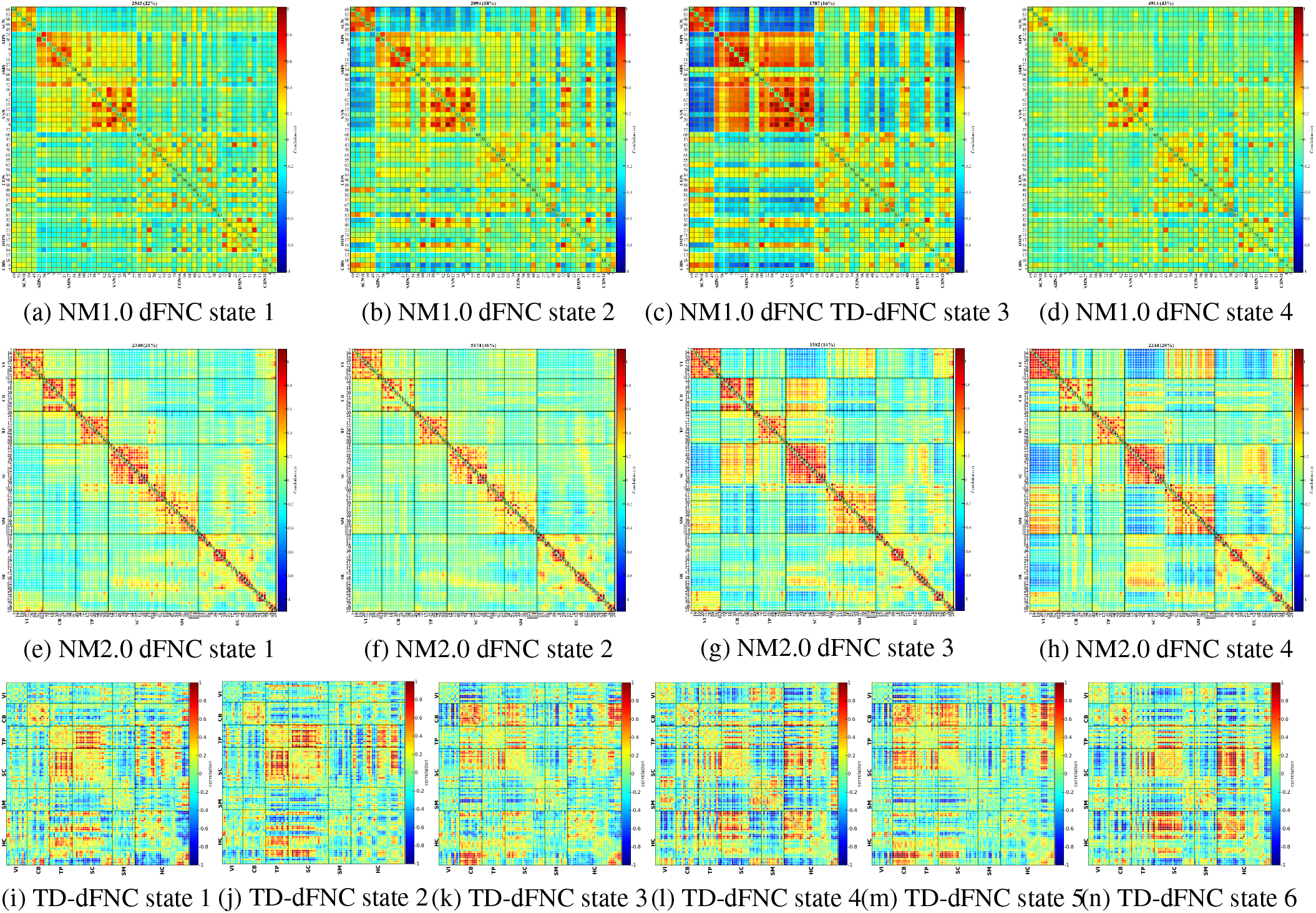
The 4 derived states from dynamic functional network connectivity (dFNC) using the Neuromark version 1.0 template (a-d), Neuromark version 2.0 template (e-h) and TD-dFNC (i-n). Each cell is the correlation value between a pair of the 53 components from the template

